# Orbitofrontal cortex to dorsal striatum circuit is critical for incubation of oxycodone craving after forced abstinence

**DOI:** 10.1101/2024.04.11.588865

**Authors:** Hongyu Lin, Adedayo Olaniran, Xiang Luo, Jessica Strauch, Megan AM Burke, Chloe L. Matheson, Xuan Li

## Abstract

Relapse is a major challenge in treating opioid addiction, including oxycodone. During abstinence, oxycodone seeking progressively increases, and we previously demonstrated a causal role of the orbitofrontal cortex (OFC) in this incubation of oxycodone craving after forced abstinence. Here, we explored critical downstream targets of OFC by focusing on dorsal striatum (DS). We first examined dorsal striatal Fos (a neuronal activity marker) expression associated with oxycodone seeking after abstinence. Using a dopamine D1 receptor (D1R) antagonist, we also tested the causal role of DS in incubated oxycodone seeking. Next, we combined fluorescence-conjugated cholera toxin subunit B (CTb-555, a retrograde tracer) with Fos to assess whether the activation of OFC**→**DS projections was associated with incubated oxycodone seeking. We then used a series of pharmacological procedures to examine the causal role of the interaction between glutamatergic projections from OFC and D1R signaling in DS in incubation of oxycodone craving. We found that dorsal striatal Fos expression in DS exhibited a time-dependent increase in parallel with incubation of oxycodone craving, and DS inactivation decreased incubated oxycodone seeking. Moreover, OFC**→**DS projections were activated during incubated oxycodone seeking, and anatomical disconnection of OFC**→**DS projections, but not unilateral inactivation of OFC or DS, decreased incubated oxycodone seeking. Lastly, contralateral disconnection of OFC**→**DS projections had no effect on oxycodone seeking on abstinence day 1. Together, these results demonstrated a causal role of OFC**→**DS projections in incubation of oxycodone craving.

## Introduction

Prescription opioids, such as oxycodone, are one of the main drivers of the ongoing opioid epidemic (Gostin et al., 2017; Stuart et al., 2018; Heins, 2019; Volkow and Blanco, 2021). A major challenge in treating opioid addiction is relapse, often triggered by drug-associated cues (O’Brien et al., 1986). We and others previously demonstrated incubation of oxycodone craving in rats, which refers to a time-dependent increase of oxycodone seeking during forced abstinence (Altshuler et al., 2021a; Altshuler et al., 2021b; Wong et al., 2022) or voluntary abstinence (Fredriksson et al., 2020; Fredriksson et al., 2021; Fredriksson et al., 2023) after oxycodone self-administration. Moreover, we identified the orbitofrontal cortex (OFC) as a critical brain region for this incubation after forced abstinence in male rats (Altshuler et al., 2021b). We found that Fos (a neuronal activity marker) expression associated with oxycodone seeking exhibits a time-dependent increase during abstinence, and OFC inactivation with muscimol+baclofen (GABAa and GABAb agonists) decreases oxycodone seeking on abstinence day 15, but not on abstinence day 1. However, circuit mechanisms underlying this incubation are largely unknown. Here, we began to explore OFC-associated circuit mechanisms by focusing on one of its downstream regions, dorsal striatum (DS). Moreover, we focused on the central to medial portion of the DS in this study. This is based on the previous anterograde tracing evidence illustrating that ventrolateral OFC, which we targeted in our previous OFC inactivation study (Altshuler et al., 2021b), projects primarily to the central to medial portion of DS (Schilman et al., 2008; Mailly et al., 2013).

Extensive literature has documented the role of DS in cocaine (Fuchs et al., 2005; Vanderschuren et al., 2005; Fuchs et al., 2006; Pacchioni et al., 2011; Murray et al., 2012), methamphetamine(Rubio et al., 2015; Li et al., 2015b; Caprioli et al., 2017), alcohol (Jeanblanc et al., 2009; Wang et al., 2010; Wang et al., 2015; Darcq et al., 2016), heroin (Rogers et al., 2008; Bossert et al., 2009) and morphine(Gao et al., 2013) seeking behaviors. To assess whether DS is a potential downstream target of OFC, we measured Fos expression associated with oxycodone seeking after abstinence and found that DS exhibited a time-dependent increase of Fos expression in parallel with incubation of oxycodone craving and associated Fos expression in OFC (Altshuler et al., 2021b). Based on this finding, we used a D1-family receptor antagonist SCH23390, which blocks drug- and cue-induced Fos expression (Valjent et al., 2000; Ciccocioppo et al., 2001; Nair et al., 2011), to test the causal role of DS in incubated oxycodone seeking, and found DS inactivation decreased incubated oxycodone seeking. These findings together set the premise for studying OFC**→**DS circuit in subsequent studies.

Growing evidence also implicates a role of OFC**→**DS projections in compulsive seeking behavior(Ahmari et al., 2013; Burguiere et al., 2013; Gremel et al., 2016; Pascoli et al., 2018) and drug-associated behaviors (Renteria et al., 2018; Hu et al., 2019; Minogianis et al., 2019; Wall et al., 2019; Cheng et al., 2021; Fredriksson et al., 2021; Pascoli et al., 2023). For example, disconnecting the OFC**→**DS circuit decreases cocaine self-administration under a progressive-ratio schedule (Minogianis et al., 2019). A recent study also demonstrated a negative correlation between OFC-DS functional connectivity and incubation of oxycodone craving after electric-barrier-imposed voluntary abstinence (Fredriksson et al., 2021). However, the role of OFC**→**DS projections in incubation of oxycodone craving has not been investigated using the forced abstinence model, the most used behavioral procedure in incubation studies (Venniro et al., 2016). Notably, emerging evidence has demonstrated distinct neural mechanisms underlying incubation of drug craving after forced abstinence versus voluntary abstinence (Venniro et al., 2020).

To study the role of OFC**→**DS projections in incubation of oxycodone craving, we first combined fluorescence-conjugated cholera toxin b (CTb-555, a retrograde tracer) with Fos and measured the activation of glutamatergic projections from OFC to DS associated with incubated oxycodone seeking. We then used a pharmacological asymmetrical disconnection approach (Bossert et al., 2012; Li et al., 2018) to examine the causal role of the interaction between OFC projections and D1R signaling in DS in incubated oxycodone seeking. In this experiment, we inactivated OFC unilaterally with muscimol+baclofen and blocked D1R signaling in either contralateral or ipsilateral DS with SCH23390. In addition, we tested whether unilateral inactivation of OFC or unilateral D1R blockade of DS alone decreased incubated oxycodone seeking and whether contralateral disconnection of OFC and DS decreased non-incubated oxycodone seeking on abstinence day 1.

## Materials and Methods

### Subjects

We used male Sprague-Dawley rats (Charles River, n=193), weighing 275-300 g prior to surgery; we maintained the rats under a reverse 12:12-h light/dark cycle with food (Teklad Global 18% protein) and water freely available. It is noted that behavioral results from Experiment 1 were published previously (Altshuler et al., 2021b). We kept the rats 4 per cage prior to surgery and then housed them individually after surgery. We performed the experiments under the protocols approved by the University of Maryland College Park Animal Care and Use Committee and in accordance with the Guide for the Care and Use of Laboratory Animals (National Institute of Health). We excluded rats due to catheter patency (n = 1), health-related issues (n = 13), cannula misplacement (n = 26), and missing brain slices (n = 1). The number of rats reported herein refers to rats included in the statistical analysis.

### Intravenous surgery

We anesthetized the rats with isoflurane (5% induction, 2-3% maintenance) and inserted silastic catheters into the rats’ jugular veins as previously described (Altshuler et al., 2021a; Altshuler et al., 2021b). We injected the rats with ketoprofen (2.5 mg/kg, s.c.) after surgery to relieve pain and inflammation; we allowed them to recover 5-7 days before oxycodone self-administration training. During the recovery and training phases, we flushed the catheters every 24-48 hours with gentamicin (Hospira; 4.8 mg/mL) diluted in sterile saline.

### Cholera toxin subunit b (CTb) injections into the DS

We injected 750 nl fluorescence-conjugated CTb (CTb-555, Invitrogen, #Catolog: C34776) unilaterally into the DS using a 10-µl, 33-gauge Nanofil syringe attached to UltraMicroPump (UMP3) with SYS-Micro4 Controller (World Precision Instruments). We counterbalanced our injections into either the left or right hemisphere. The concentration of CTb-555 was 4 µg/µl, dissolved in phosphate-buffered saline (PBS) (Mandelbaum et al., 2019). The CTb-555 was delivered over 5 min with the needle left in place for an additional 5 min. Based on our previous study (Li et al., 2015b; Li et al., 2018), we used the following coordinates for DS: anteroposterior (AP), +1.6 mm; mediolateral (ML), ±2.4 mm; dorsoventral (DV), -5.4 mm.

### Cannula implantations

We implanted guide cannulas (23 gauge; Plastics One) 1.0 mm above the OFC or DS with the nose bar set at -3.3 mm. Based on our previous study (Altshuler et al., 2021b), we used the following coordinates for OFC: AP, +3.7 mm; ML, ±3.4 (10° angle); DV, -3.9 mm. We used the following coordinates for DS in bilateral DS inactivation, contralateral disconnection, and unilateral DS inactivation studies: AP, +1.6 mm; ML, ±3.0 mm (6° angle); DV, -4.4 mm, based on previous studies (Li et al., 2015b; Li et al., 2018). For the ipsilateral inactivation study, we used the following coordinate coordinates for DS: AP, +1.6 mm; ML, ±1.0 mm (6° angle); DV, -5.0mm, and we inserted the cannula at a 6° angle from the contralateral hemisphere. We anchored the cannulas to the skull with jeweler’s screws and dental cement.

### Intracranial injections

We dissolved muscimol + baclofen (Tocris Bioscience) or SCH23390 (Tocris Bioscience) in sterile saline and injected the drugs 15 min before starting the seeking test. The doses of muscimol+baclofen (50 + 50 ng in 0.5µl/side), SCH23390 (0.75 µg in 0.5 µl/side), and drug pretreatment time were based on our previous studies (Li et al., 2015b; Altshuler et al., 2021b). For intracranial injection, we used 10-µl Hamilton syringes attached to a syringe pump (Harvard Apparatus). The syringes were connected to the 30-gauge injectors via polyethylene-50 tubing. The injectors extended 1.0 mm below the tips of the guide cannulas for the OFC or DS. The drugs were delivered at a rate of 0.5 µl/min with the injectors in place for an additional minute to allow diffusion. After the seeking test, rats’ brains were extracted and stored in 10% formalin. Two days later, we sliced the rat brains (50 µm sections) using a Leica cryostat and stained the sections with cresyl violet. Finally, cannula placements were verified using a Nikon DS-Fi3 camera attached to an inverted Nikon Eclipse Ti2 Series microscope.

### Apparatus

We trained the rats in self-administration chambers located inside sound-attenuating cabinets and controlled by a Med Associates (Georgia, VT) system. Each chamber has two levers located 8-9 cm above the floor. During self-administration training, presses on the retractable (active) lever activated the infusion pump (which delivered an oxycodone infusion); presses on the stationary (inactive) lever were not reinforced with the drug. For oxycodone intravenous infusions, each rat’s catheter was connected to a liquid swivel (Instech) via polyethylene-50 tubing, protected by a metal spring. We then attached the liquid swivel to a 20-ml syringe via polyethylene-50 tubing and to a 22-gauge modified needle (Plastics One, VA).

### Oxycodone self-administration training

We used a training procedure as described in our previous studies (Altshuler et al., 2021a; Altshuler et al., 2021b; Olaniran et al., 2023). Briefly, rats were trained to self-administer oxycodone for 6 h per day (six 1-h sessions with 10 minutes off in between each session), under a fixed-ratio-1 (FR1) with 20-s timeout reinforcement schedule, for 10 sessions over an 11-day period (off day between 5^th^ and 6^th^ day). The oxycodone was delivered at a dose of 0.1 mg/kg/infusion over 3.5 s (0.10 ml/infusion). We used Brevital (3 to 4 mg/kg) to check catheter patency for low responders during the training.

The daily training sessions started at the onset of the dark cycle and began with the extension of the active lever and the illumination of the red house light. The house light remained on for the duration of each 1-h session for a total of six 1-h sessions. During training, active lever presses led to the delivery of an oxycodone infusion and a compound 5-s tone-light cue (the tone and light modules were located above the active lever). During the 20-s timeout, we recorded the non-reinforced lever presses. We set 15 infusions as the maximum for each 1-h session to prevent overdose. The red house light was turned off, and the active lever retracted after the rats received the maximum number of infusions or at the end of the 6-h session. For all experiments, we counterbalanced their oxycodone between experimental groups based on intake during oxycodone self-administration.

### Abstinence phase

During the abstinence phase, we housed the rats individually in the animal facility and handled them 2- 3 times per week.

### Oxycodone seeking test (Seeking test)

We conducted all seeking tests (2 h) immediately after the onset of the dark cycle. The sessions began with the extension of the active lever and the illumination of the red house light, which remained on for the duration of the session. Active lever presses during testing [the operational measure of drug seeking in incubation of craving studies (Lu et al., 2004; Pickens et al., 2011) resulted in contingent presentations of the tone-light cue, previously paired with oxycodone infusions, but did not result in drug infusions. Inactive lever presses were used as a measure of non-specific activity and/or response generalization (Lu et al., 2004).

### Food self-administration

To facilitate the acquisition of food self-administration, we gave 1-h magazine training before the operant training during the first two training days. During magazine training, we presented the food pellets to the rats every 5 min; pellet delivery was paired with a 5-s light cue. Then we trained rats to self-administer food pellets (Test Diet, #1811155) for 1 h per day during the middle of their dark cycle, under an FR1 20-s timeout reinforcement schedule; pellet delivery was paired with 5-s light cue. To increase rats’ motivation to press for food pellets, we restricted their chow in home cage to 20 g/day and fed them after they finished the food self-administration session.

### Fos Immunohistochemistry (IHC)

Immediately after the seeking tests on abstinence day 15, we anesthetized the rats with isoflurane and perfused them transcardially with ∼100 ml of 0.1 M PBS followed by 400 ml of 4% paraformaldehyde (PFA) in PBS. We extracted the brains, which were then postfixed in 4% PFA for 2 h before being transferred to 30% sucrose in PBS for 48 h at 4°C. After freezing the brains on dry ice, we cut serial coronal sections (40 µm) using a Leica Microsystems cryostat and preserved the sections in cryoprotectant (20% glycerol and 2% DMSO in PBS, pH 7.4).

We processed a one-in-five series of sections from the OFC from each rat for immunochemical detection of Fos. We repeatedly rinsed free-floating sections in PBS (3 x 10 min) and incubated them for 1 h in 10% normal horse serum (NHS) in PBS with 0.5% Triton X-100 (PBS-Tx). All sections were incubated overnight at 4⁰ C with rabbit anti-c-Fos primary antibody (5348, 1:2000; Cell Signaling Technology, RRID: AB_10557109) diluted in 2% NHS in PBS-Tx.

For Experiment 1, the sections were washed with PBS (3 x 10 min) and incubated with biotinylated goat anti-rabbit secondary antibody (1:600, Vector Laboratories, Cat# BA-1000, RRID:AB_2313606), diluted in 1% NGS in PBS with Triton X-100, for 2 h at room temperature. Next, we washed the sections with PBS (3 x 10 min washes) and incubated the sections with avidin-biotin-peroxidase complex (ABC, ABC Elite Kit, #PK6100, Vector Laboratories) for 1 h at room temperature. We then washed the sections with PBS (3 x 10 min) and developed the sections in 3,3’-Diaminobenzidine (DAB) for 100 s. The sections were washed in PBS (4 x 5 min) and mounted on glass slides (Fisherbrand™ Superfrost™ Plus Microscope Slides, Cat #12-550-15). Once dried, the slides were dehydrated in a series of ethanol (30%, 60%, 90%, 95%, 100%, 100%) and cleaned with Citrisolv (Fisher Scientific). We then cover-slipped slides with Permount (Fisher Scientific).

For Experiment 3, we washed off unbound primary antibody with PBS and incubated sections with donkey anti-rabbit Alexa-Fluor-647 (31573, 1:500; Invitrogen, RRID: AB_2536183) for 2-4 h in 2% NHS in PBS-Tx. After rinsing sections in PBS, we mounted the sections onto glass slides (Basix^TM^ Adhesion Microscope Slides, Cat #23-888-115). These sections were then air dried, and coverslipped with Fluormount G (Electron Microscopy Sciences).

### Image acquisition and neuronal quantification

For Experiment 1, we digitally captured bright-field images of Fos immunoreactive (IR) cells in DS using a Nikon DS-Fi3 camera attached to an inverted Nikon Eclipse Ti2 Series microscope. We captured and analyzed the images using NIS-Elements (Nikon, 5.20.00) at 10X magnification in a blind manner. We used an automatic counting method to quantify Fos-IR cells (inter-rater reliability between H.L and MMD r = 0.98, p < 0.05). For each rat, we analyzed two brain sections (4 hemispheres) in DS using the following bregma levels: +1.6 to +1.2mm.

For Experiment 3, we digitally captured dark-field images of CTb-555 and Fos immunoreactive (IR) cells in OFC using a Hamamatsu Flash 4.0 LT Plus camera attached to the Nikon Ti2 microscope. We analyzed the images at 10 X magnification in a blind manner, using the General Analysis Package of NIS-Elements (version 5.20.00); inter-rater reliability between H.L. and Xiang.L. for CTb-555 is r =0.73, p< 0.05; for CTb+Fos is r =0.85, p< 0.05). For each rat, we quantified cells in both ipsilateral and contralateral hemisphere to the CTb-555 injection. We analyzed four sections/rat using the following bregma range for OFC: +3.72 to +3.0 mm.

### Experimental procedures

#### Experiment 1: dorsal striatal Fos expression associated with oxycodone seeking after abstinence

We performed Fos IHC in DS from 4 groups of rats (n=23) that received training for oxycodone self-administration as described above. On either abstinence day 1 or 15, one group of rats were tested for oxycodone seeking (Day 1: n = 6; Day 15: n = 6) and the other group of rats served as No-test groups (Day 1: n = 5; Day 15: n = 6). All rats were perfused either directly from the home cage (No-test groups) or immediately after the 2-h seeking test session (Test groups), for subsequent IHC assays. The behavioral results from this experiment were previously published (Altshuler et al., 2021b). Here, we measured Fos expression in three subregions of DS: dorsomedial striatum (DMS), dorsocentral striatum (DCS), and dorsolateral striatum (DLS). This is based on both the topographical organization of OFC**→**DS projections (Schilman et al., 2008; Mailly et al., 2013) and distinct roles of DMS and DLS in drug-seeking behaviors (Vanderschuren et al., 2005; Bossert et al., 2009; Wang et al., 2010; Corbit et al., 2012; Rubio et al., 2015; Caprioli et al., 2017).

#### Experiment 2: the causal role of DS in oxycodone seeking on abstinence day 15

We performed intravenous surgeries on two groups of rats (n = 10) and implanted bilateral guide cannulas 1.0 mm above the DS. We then trained all rats to self-administer oxycodone as described above. On abstinence day 15, we injected the rats with vehicle (n=5) or D1-family receptor antagonist (SCH23390, 0.75 µg/0.5 µl/side, n=5) bilaterally into DS, 15 min before the 2-h seeking tests.

#### Experiment 3: activation of OFC**→**DS projections associated with oxycodone seeking on abstinence day 15

We performed intravenous surgeries and CTb-555 injections into the DS on two groups of rats (n = 11) and trained them for oxycodone self-administration. On abstinence day 1, we tested one group of rats for oxycodone seeking in a 2-h session, and this group later served as the No-test group (n=4) on abstinence day 15. On abstinence day 15, we tested the second group of rats for oxycodone seeking (Seeking test, n=7). Immediately after the seeking test, we anesthetized the rats, perfused them, and extracted the brains of the rats from the Seeking-test group. During the same time, we perfused and extracted the brains of the rats from the No-test group, which we brought to the perfusion station directly from their home cages. We measured CTb and CTb+Fos double-labeled cells in the OFC.

#### Experiment 4: effect of contralateral and ipsilateral disconnection of OFC and DS on oxycodone seeking on abstinence day 15

We performed intravenous surgeries on 4 groups of rats (n = 31) and implanted each rat with two unilateral cannulas 1 mm above OFC or DS. The two unilateral cannulas were either in the opposite (contralateral) or the same hemisphere (ipsilateral). We counterbalanced the left and right hemispheres for cannula implantation. After training all rats for oxycodone self-administration, we performed contralateral disconnection of OFC**→**DS projections by injecting muscimol+baclofen (50+50 ng//0.5 µl/side) into OFC and SCH23390 (0.75 µg/0.5 µl/side) to the contralateral DS on abstinence day 15 (n=8). For ipsilateral disconnection, we injected muscimol+baclofen (50+50 ng/0.5 µl/side) into OFC and SCH23390 (0.75 µg/0.5 µl/side) to the ipsilateral DS on abstinence day 15 (n=6). Vehicle rats received saline injections in both OFC and DS (Contralateral Vehicle: n=10; Ipsilateral Vehicle: n=7). We tested all rats for oxycodone seeking 15 min after intracranial injections.

To examine whether ipsilateral disconnection of OFC and DS caused motor deficits, we used a within-subject design and trained rats from the vehicle group of the ipsilateral disconnection experiment to self-administer palatable food pellets (n=6) for 4 d (1 h/d), as described above. We then injected the rats with either saline to both OFC and ipsilateral DS, or muscimol+baclofen (50+50 ng/0.5 µl/side) into OFC and SCH23390 (0.75 µg/0.5 µl/side) into ipsilateral DS, 15 min before the 1-h food self-administration session on 5th and 7th day. We re-trained rats on the 6^th^ day. The order of the vehicle and drug injections was counterbalanced across testing days.

#### Experiment 5: effect of unilateral inactivation of OFC or DS on oxycodone seeking on abstinence day 15

We performed intravenous surgeries on four groups of rats (total n = 31) and implanted each rat with one unilateral cannula 1 mm above the OFC or DS. We counterbalanced the left and right hemispheres for cannula implantation. We then trained the rats for oxycodone self-administration. On abstinence day 15, we injected either muscimol+baclofen (50+50 ng/0.5 µl/side) into OFC (n=8) or SCH23390 (0.75 µg/0.5 µl/side) into DS (n=9). Vehicle rats received saline injections (OFC vehicle: n=7; DS vehicle: n=7). All rats were tested for oxycodone seeking 15 min after the intracranial injections.

#### Experiment 6: effect of contralateral disconnection of OFC and DS on oxycodone seeking on abstinence day 1

We performed intravenous surgeries on 2 groups of rats (n = 15) and implanted each rat with one cannula 1 mm above OFC and one cannula 1 mm above the contralateral DS. We counterbalanced the left and right hemispheres for cannula implantation. After training all rats for oxycodone self-administration, we performed contralateral disconnection of OFC**→**DS projections by injecting muscimol+baclofen (50+50 ng/0.5 µl/side) into OFC or SCH23390 (0.75 µg/0.5 µl/side) to the contralateral DS on abstinence day 1 (n=7). Vehicle rats received saline injections in both OFC and DS (n=8).

To examine whether contralateral disconnection of OFC and DS caused motor deficits, we used a within-subject design and trained rats from the vehicle group of this experiment to self-administer palatable food pellets (n=5) for 5 d (1 h/d), as described above. We then injected the rats with either saline to both OFC and DS, or muscimol+baclofen (50+50 ng/0.5 µl/side) into OFC or SCH23390 (0.75 µg/0.5 µl/side) into contralateral DS, 15 min before the 1-h food self-administration session on the 5th and 11th day. We re-trained rats from the 6^th^ to 10^th^ day. The order of the vehicle and drug injections was counterbalanced across testing days.

### Statistical analysis

We analyze the data with SPSS (version 27) using repeated measures, mixed-factorial ANOVAs (General Linear Model), or t-tests. We used univariate ANOVAs to perform post hoc tests following significant interaction or main effects. We only report post hoc effects that are critical for data interpretation. For the repeated measures analyses of the training data, we replaced 51 outlier values of active lever presses and 37 outlier values of inactive lever presses with the group mean for a given training day. We defined outliers as three median absolute deviations (MADs) above the group median(Leys et al., 2013), and we only replaced one outlier (the highest value above the threshold) for each training day. For rats that received fewer than 20 daily fusions on average (n = 7) during the 10 training days, we extended their training for an additional 1–5 days, and we used the last 10 training days for the data analysis. We indicate the between- and within-subject factors of the different analyses in the Results. All statistical comparisons are listed in Tables S1 and S2.

## Results

### Oxycodone self-administration (Experiment 1-5)

Rats in all experiments demonstrated escalation of oxycodone self-administration and showed a strong preference for the oxycodone-associated active lever over the inactive lever during the training phase (Figs. 2B, 3C, 4B, 4C, 5B, 5C and 6B). Detailed statistical information is provided in Table S1.

**Figure 1.**
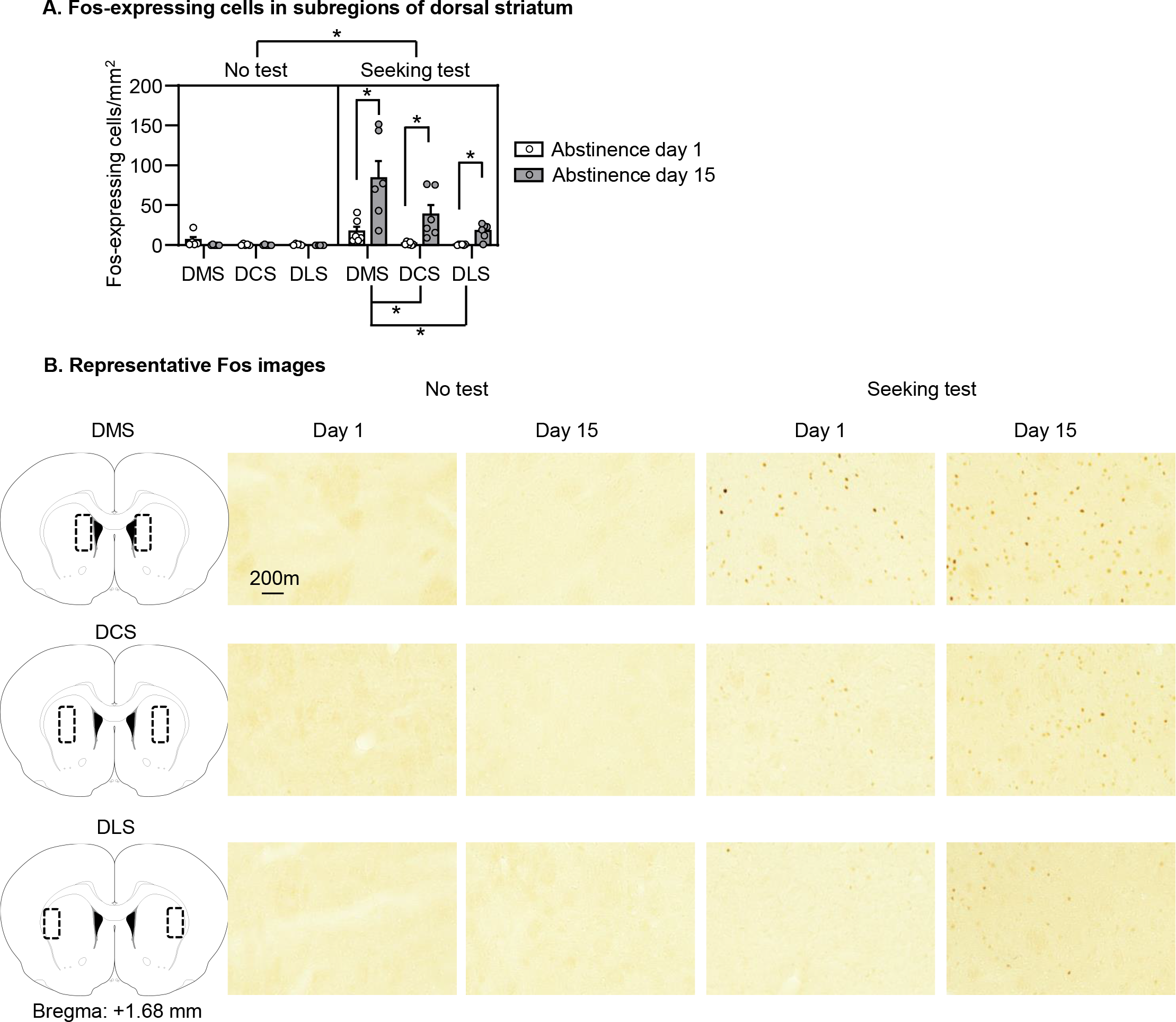
*DS exhibited time-dependent increases of Fos expression associated with oxycodone seeking after abstinence.* **(A)** Data are the mean ± SEM of Fos/mm^2^ in dorsomedial striatum (DMS), dorsocentral striatum (DCS), and dorsolateral striatum (DLS) after the seeking test on either abstinence day 1 or 15. *Different between two groups as marked, p<0.05. Day-1 No test: n=5; Day 1 Seeking test: n=6; Day-15 No test: n=6; Day 15 Seeking test: n=6 **(B)** Representative images of Fos labeling in DMS, DCS, and DLS.

**Figure 2.**
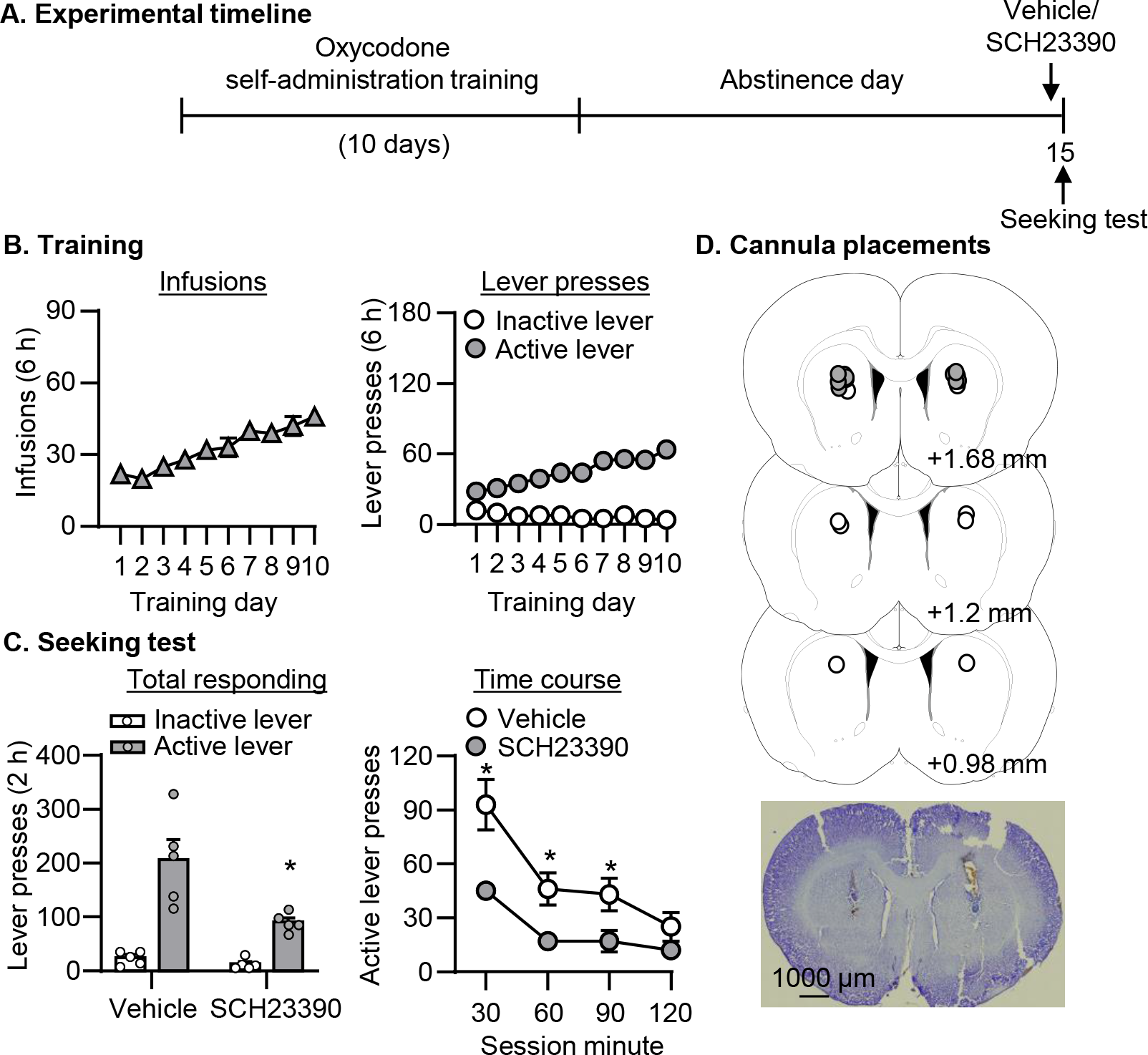
*Bilateral SCH23390 injections into DS decreased oxycodone seeking on abstinence day 15 (Experiment 2).* **(A)** Timeline of the experiment. **(B)** Oxycodone self-administration. Data are mean ± SEM number of oxycodone (0.1 mg/kg/infusion) infusions or lever presses during the 10 6-h daily self- administration sessions (total n = 10). **(C)** Seeking test. Data are mean ± SEM of lever presses on the previously active lever and on the inactive lever during the seeking test sessions. *Different from Vehicle, p<0.05, Vehicle: n = 5, SCH23390: n = 5. **(D)** Approximate placement [mm from Bregma (Paxinos and Watson, 2005)] of injection tips (vehicle: open circles; SCH23390: closed circles), and representative cannula placements.

**Figure 3.**
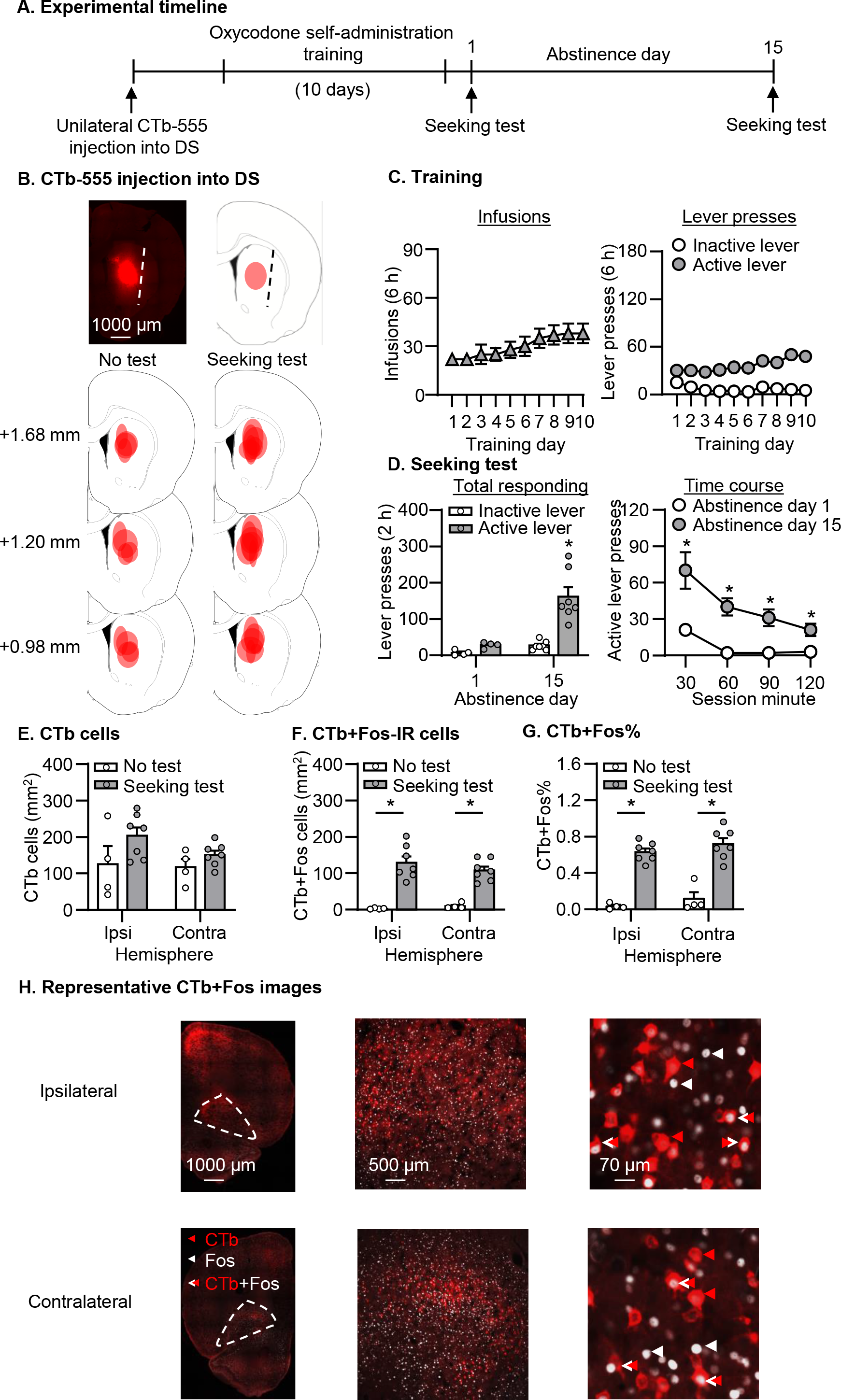
*Incubated oxycodone seeking is associated with activation of OFC**→**DS projections (Experiment 3).* **(A)** Timeline of the experiment. **(B)** Representative plots of the spread of CTb-555 injections in No-test and Seeking-test group. **(C)** Oxycodone self-administration. Data are mean ± SEM number of oxycodone (0.1 mg/kg/infusion) infusions or lever presses during the 10 6-h daily self-administration sessions (total n = 11). **(D)** Seeking tests on abstinence days 1 and 15. Data are mean ± SEM of lever presses on the previously active lever and on the inactive lever during the seeking test sessions. *Different from Vehicle, p<0.05, No test: n = 4, Test: n = 7. **(E-G)** Data are the mean ± SEM of CTb/mm^2^, CTb+Fos/mm^2^ and Fos% in OFC after the seeking test on abstinence day 15. *Different from No test, p<0.05, No test: n = 4, Test: n = 7. **(H)** Representative images of CTb and Fos labeling in OFC.

**Figure 4.**
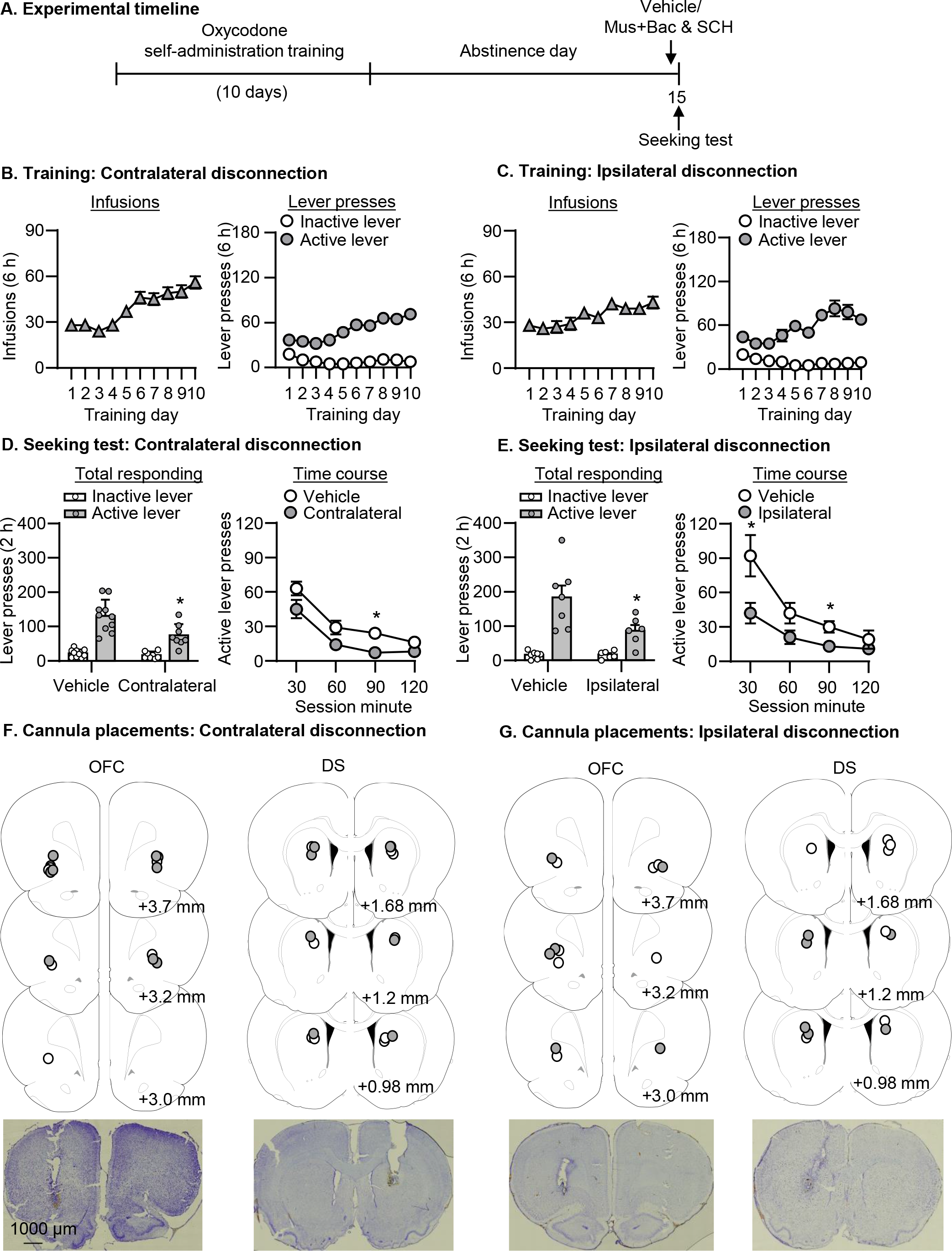
Contralateral and ipsilateral disconnection of OFC and DS decreased *oxycodone seeking on abstinence day 15 (Experiment 4).* **(A)** Timeline of the experiment. **(B-C)** Oxycodone self-administration. Data are mean ± SEM number of oxycodone (0.1 mg/kg/infusion) infusions or lever presses during the 10 6-h daily self-administration sessions (contralateral: total n = 18; ipsilateral: total n = 13). **(D-E)** Seeking tests. Data are mean ± SEM of lever presses on the previously active lever and on the inactive lever during the seeking test sessions. *Different from Vehicle, p < 0.05, Contralateral: Vehicle, n = 10; Contralateral, n = 8. Ipsilateral: Vehicle, n = 7; Ipsilateral, n = 6. **(F-G)** Approximate placement [mm from Bregma (Paxinos and Watson, 2005)] of injection tips (vehicle: open circles; Muscimol+baclofen or SCH23390: closed circles), and representative cannula placements.

#### Experiment 1: dorsal striatal Fos expression associated with oxycodone seeking after abstinence (Fig. 1)

As a first step to examine whether DS is a potential downstream target of OFC during incubation of oxycodone craving, we measured Fos expression in DMS, DCS and DLS associated with oxycodone seeking on either abstinence day 1 or day 15. We found that all three DS subregions exhibited time-dependent increases of Fos expression in parallel with incubation of oxycodone craving and OFC activation observed in our previous study (Altshuler et al., 2021b) . On both abstinence days 1 and 15, we found oxycodone seeking was associated with a higher number of Fos-IR cells in DMS than DCS and DLS.

We first analyzed the Fos data using a three-way repeated ANOVA with the between-subject factors of Abstinence day (Day 1, Day 15) and test condition (No test, Seeking test), and the within-subject factor of Subregion (DMS, DCS, DLS). We found a significant triple interaction among these three factors (F2,38 = 5.415, p=0.009). We followed up on this interaction by dividing the 3-way ANOVA into the following two-way ANOVA analyses. First, we analyzed the Fos data within each DS subregion using the between-subject factors of Abstinence Day (Day 1, Day 15) and test condition (No test, Seeking test). We found a significant interaction between these two factors across all three DS subregions (DMS: F1,19 = 9.066, p=0.007; DCS: F1,19 = 8.104, p=0.01; DLS: F1,19 = 18.214, p<0.001). *Post hoc* analyses showed that there were significantly higher numbers of Fos-IR cells in Day-15 Test group than Day-1 Test group across three DS subregions (p<0.05). Next, we analyzed the Fos data for each test condition using the between-subject factor of Abstinence Day (Day 1, Day 15) and the within-subject factor of Subregion (DMS, DCS, DLS). We found a significant interaction between Abstinence Day and Subregion for Seeking Test condition (F2,20 = 4.979, p=0.018), but not No-test condition (p>0.05). *Post hoc* analyses showed that there were significantly higher numbers of Fos-IR cells in DMS than DCS and DLS on both abstinence days 1 and 15 (p<0.05).

These findings support the hypothesis that DS is a potential downstream target of OFC during incubation of oxycodone craving. Based on this finding, we tested the causal role of DS in incubated oxycodone seeking in Experiment 2. We also observed that oxycodone seeking was associated with more Fos expression in DMS than in DCS and DLS. Together with anatomical evidence from previous studies (Schilman et al., 2008; Mailly et al., 2013) regarding the topographical organization of OFC**→**DS projection, we focused on the DMS and DCS (together referred as DS) for subsequent studies.

#### Experiment 2: the causal role of DS in oxycodone seeking on abstinence day 15 (Fig. 2)

In this experiment, we used a D1-family receptor antagonist SCH23390, which blocks drug- and cue-induced Fos expression (Valjent et al., 2000; Ciccocioppo et al., 2001; Nair et al., 2011), to test the causal role of DS in incubated oxycodone seeking. We found that bilateral injections of SCH23390 into DS decreased oxycodone seeking on abstinence day 15 (Fig. 2C). The analysis, which included the between-subject factor of Treatment (Vehicle, SCH23390) and within-subject factor of Lever (Active lever, Inactive lever), showed a significant interaction between Treatment and Lever (F1,8 = 9.528, p=0.015). *Post hoc* analyses showed that there were significantly lower active lever presses in rats with SCH23390 injections than rats with vehicle injections (p<0.05). To examine the effect of SCH23390 on within-session extinction learning, we also analyzed the time course data of active lever presses by using the between-subject factor of Treatment (Vehicle, SCH23390) and the within-subject factor of Session minute (30, 60, 90, 120). We found a significant interaction between these two factors (F3,24 = 4.477, p=0.012). It is noted that the dose of SCH23390 used here does not cause motor deficits, as our previous study demonstrated that SCH23390 injections into DS at the same dose have no effect on ongoing food self-administration (Li et al., 2015b). Together with Experiment 1, these data demonstrated that DS played a causal role in incubated oxycodone seeking and set premises for the subsequent experiments focusing on the role of OFC**→**DS projections in incubation of oxycodone craving.

#### Experiment 3: activation of OFC**→**DS projections associated with oxycodone seeking on abstinence day 15 (Fig. 3)

In Experiment 3, we asked whether activation of OFC**→**DS projections is associated with incubated oxycodone seeking on abstinence day 15. To answer this question, we used Fos in combination with fluorescence-conjugated retrograde tracer (CTb-555, injected into DS) and measured CTb+Fos double labeled cells in OFC after the seeking test on abstinence day 15.

##### Seeking test (*Fig. 3D*)

Oxycodone seeking was higher on abstinence day 15 than on day 1 (Fig. 3D), demonstrating “incubation of oxycodone craving” under our experimental condition. We analyzed the data with the between-subject factor of Abstinence day (Day 1, Day 15) and within-subjects factor of Lever (Active lever, Inactive lever). We observed a significant interaction between Abstinence day and Lever (F1,9 = 12.128, p=0.007). *Post hoc* analyses indicated that active lever presses were significantly higher on abstinence day 15 than on day 1 (p<0.05). We analyzed the time course data of active lever presses by using the between-subject factor of Abstinence day (Day 1, Day 15) and the within-subject factor of Session minute (30, 60, 90, 120). We found a main effect of Abstinence day (F1,9 = 14.074, p=0.005) and Session minute (F3,27 = 8.953, p<0.001), but no significant interaction between these two factors (p>0.05).

##### CTb and CTb+Fos data (*Fig. 3E-G*)

There was no difference in the total number of CTb cells between No test and Seeking test groups or between hemispheres (Fig. 3E). The number of CTb+Fos cells (Fig. 3F) and the percentage of Fos (out of the total number CTb cells, Fig. 3G) in both ipsilateral and contralateral OFC significantly increased after the oxycodone seeking test. The analysis of the number of CTb cells, which included the between-subject factor of test condition (No test, Seeking test) and within-subject factor of Hemisphere (Ipsilateral, Contralateral) showed no main effects of either factor, or interaction between two factors (p>0.05). The analysis of the number of CTb+Fos cells and the percentage of Fos, which included the between-subject factor of test condition (No test, Seeking test) and within-subject factor of Hemisphere (Ipsilateral, Contralateral) showed a significant main effect of test condition (Fos: F1,9 = 97.187, p<0.001; CTb+Fos: F1,9 = 36.734, p<0.001), but no main effect of Hemisphere or interaction between these two factors (p>0.05).

These data demonstrated that activation of both ipsilateral and contralateral OFC**→**DS projections was associated with incubated oxycodone seeking. Based on this finding, we examined the causal role of OFC**→**DS projections in incubation of oxycodone craving.

#### Experiment 4: effect of contralateral and ipsilateral disconnection of OFC and DS on oxycodone seeking on abstinence day 15 (Fig. 4)

Our recent publication demonstrated that inactivating bilateral OFC with muscimol+baclofen decreased incubated oxycodone seeking (Altshuler et al., 2021b). Together with results from Experiments 2 and 3, we aimed to examine the causal role of the interaction between activated OFC**→**DS projection and D1R signaling in DS in incubated oxycodone seeking. We used an asymmetric pharmacological disconnection procedure, in which we inhibited glutamatergic neurons in OFC unilaterally using muscimol+baclofen and D1Rs in either contralateral or ipsilateral DS using SCH23390.

We found that both contralateral and ipsilateral disconnection of OFC and DS decreased oxycodone seeking on abstinence day 15 (Figs. 4D and 4E). We analyzed the total responding using the between-subject factor of Treatment [Vehicle, Contralateral (or Ipsilateral) disconnection] and within-subject factor of Lever (Active lever, Inactive lever), and we observed a significant interaction between Treatment and Lever (Contralateral disconnection: F1,16 = 9.511, p = 0.007; Ipsilateral disconnection: F1,11 = 6.248, p = 0.03). *Post hoc* analyses indicated that there were significantly lower active lever presses in both contralateral and ipsilateral disconnection groups compared with their respective control groups (p<0.05). We also analyzed the time course data of active lever presses by using the between-subjects factor of Treatment [Vehicle, Contralateral (or Ipsilateral) disconnection] and the within-subject factor of Session minute (30, 60, 90, 120). In contralateral disconnection, we found significant main effects of Treatment (F1,16 = 8.362, p=0.011) and Session minute (F3,48 = 53.141, p<0.001), but no interaction between these two factors. In ipsilateral disconnection, we found a significant interaction between Treatment and Session minute (F3,33 = 3.62, p=0.023). Together, these data demonstrated a critical role of OFC**→**DS projections in incubated oxycodone seeking.

#### Experiment 5: effect of unilateral inactivation of OFC or DS on oxycodone seeking on abstinence day 15 (Fig. 5)

The findings in Experiment 4 suggested the possibility that disrupting unilateral neural activity in OFC or unilateral D1R signaling in DS may be sufficient to interfere with incubated oxycodone seeking, which could explain the similar inhibitory effects after either contralateral or ipsilateral disconnection of OFC and DS. To explore this possibility, we examined the effect of unilateral inactivation of OFC or DS on oxycodone seeking on abstinence day 15. We found that unilateral inactivation of neither OFC by muscimol+baclofen nor DS by SCH23390 impacted incubated oxycodone seeking (Figs. 5D and 5E). We analyzed the total responding using the between-subject factor of Treatment [Vehicle, Muscimol+Baclofen (or SCH2330)] and within-subject factor of Lever (Active lever, Inactive lever). We observed significant main effects of Lever (unilateral OFC: F1,13 = 70.485, p<0.001; unilateral DS: F1,14 = 27.735, p<0.001) but no effect of Treatment or interaction between these factors (p > 0.05). We also analyzed the time course data of active lever presses by using the between-subjects factor of Treatment [Vehicle, Muscimol+Baclofen (or SCH2330)] and the within-subject factor of Session minute (30, 60, 90, 120). We found significant main effects of Session minute (OFC: F3,39 = 34.485, p<0.001; DS: F3,42 = 32.689, p<0.001), but no main effects of Treatment or interaction between these two factors (p > 0.05).

**Figure 5.**
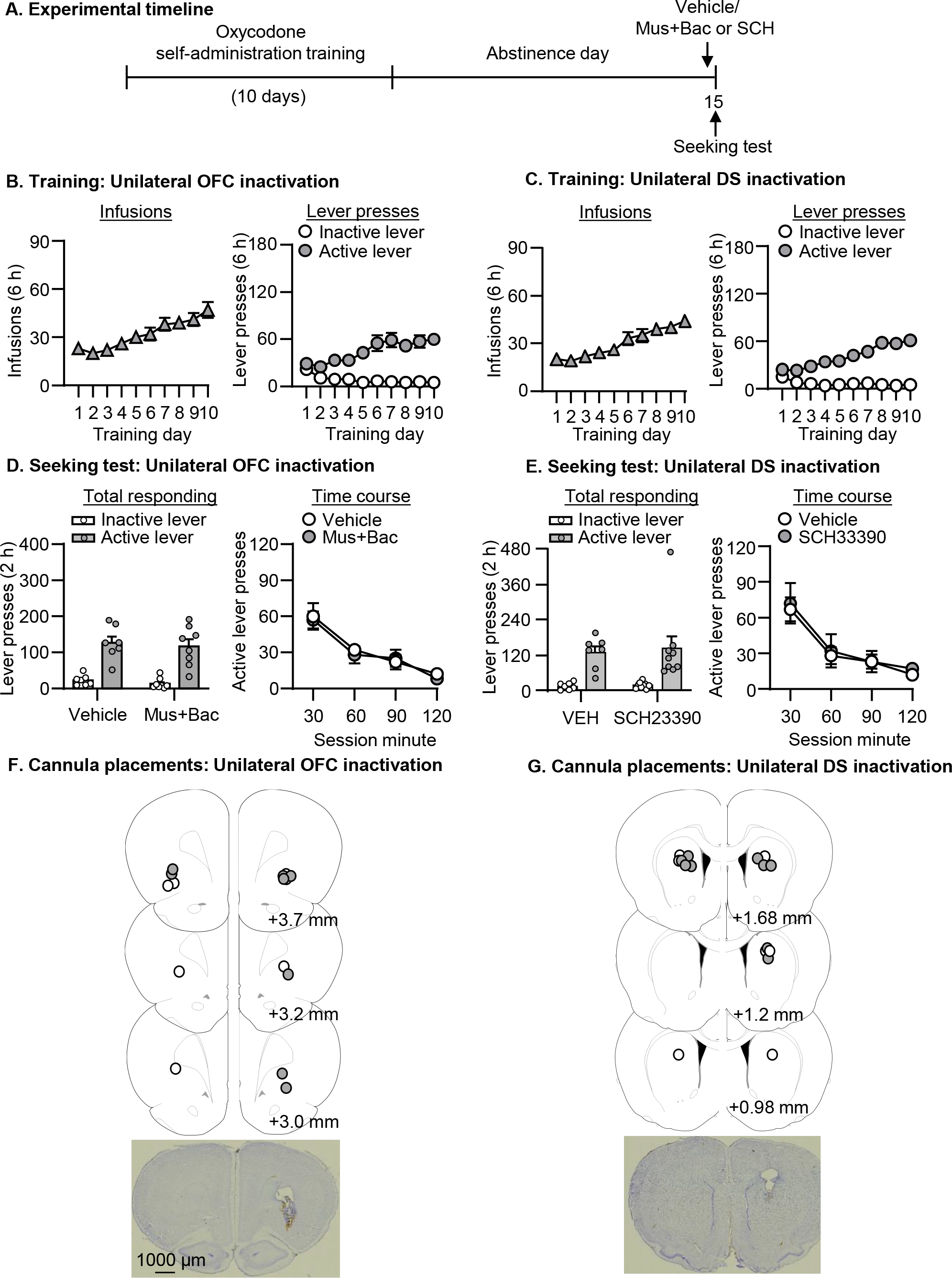
*Unilateral inactivation of OFC or DS had no effect on oxycodone seeking on abstinence day 15 (Experiment 5).* **(A)** Timeline of the experiment. **(B-C)** Oxycodone self-administration. Data are mean ± SEM number of oxycodone (0.1 mg/kg/infusion) infusions or lever presses during the 10 6-h daily self- administration sessions (OFC: total n = 15; DS: total n = 16). **(D-E)** Seeking tests. Data are mean ± SEM of lever presses on the previously active lever and on the inactive lever during the seeking test sessions. OFC: Vehicle, n = 7; Mus+Bac, n = 8. DS: Vehicle, n = 7; SCH23390, n = 9. **(F-G)** Approximate placement [mm from Bregma (Paxinos and Watson, 2005)] of injection tips (vehicle: open circles; Muscimol+baclofen or SCH23390: closed circles), and representative cannula placements.

#### Experiment 6: effect of contralateral disconnection of OFC and DS on oxycodone seeking on abstinence day 1 (*Fig. 6*)

In this experiment, we asked whether the critical role of OFC**→**DS in incubated oxycodone seeking (observed in Experiment 4) also generalized to non-incubated oxycodone seeking on abstinence day 1. To answer this question, we used the pharmacological disconnection procedure as described in Experiment 3 and examined the effect of contralateral disconnection of OFC and DS on oxycodone seeking on abstinence day 1. We found that contralateral disconnection of OFC and DS had no effect on non-incubated oxycodone seeking on abstinence day 1 (Fig. 6C). We analyzed the total responding using the between-subject factor of Treatment (Vehicle, Contralateral disconnection) and within-subject factor of Lever (Active lever, Inactive lever). We observed main effects of Lever (F1,13 = 27.377, p<0.001), but no main effect of Treatment or interaction between these factors (p > 0.05). We also analyzed the time course data of active lever presses by using the between-subjects factor of Treatment (Vehicle, Contralateral disconnection) and the within-subject factor of Session minute (30, 60, 90, 120). We found significant main effects of Session minute (F3,39 = 15.004, p<0.001), but no main effects of Treatment or interaction between these two factors (p > 0.05).

**Figure 6.**
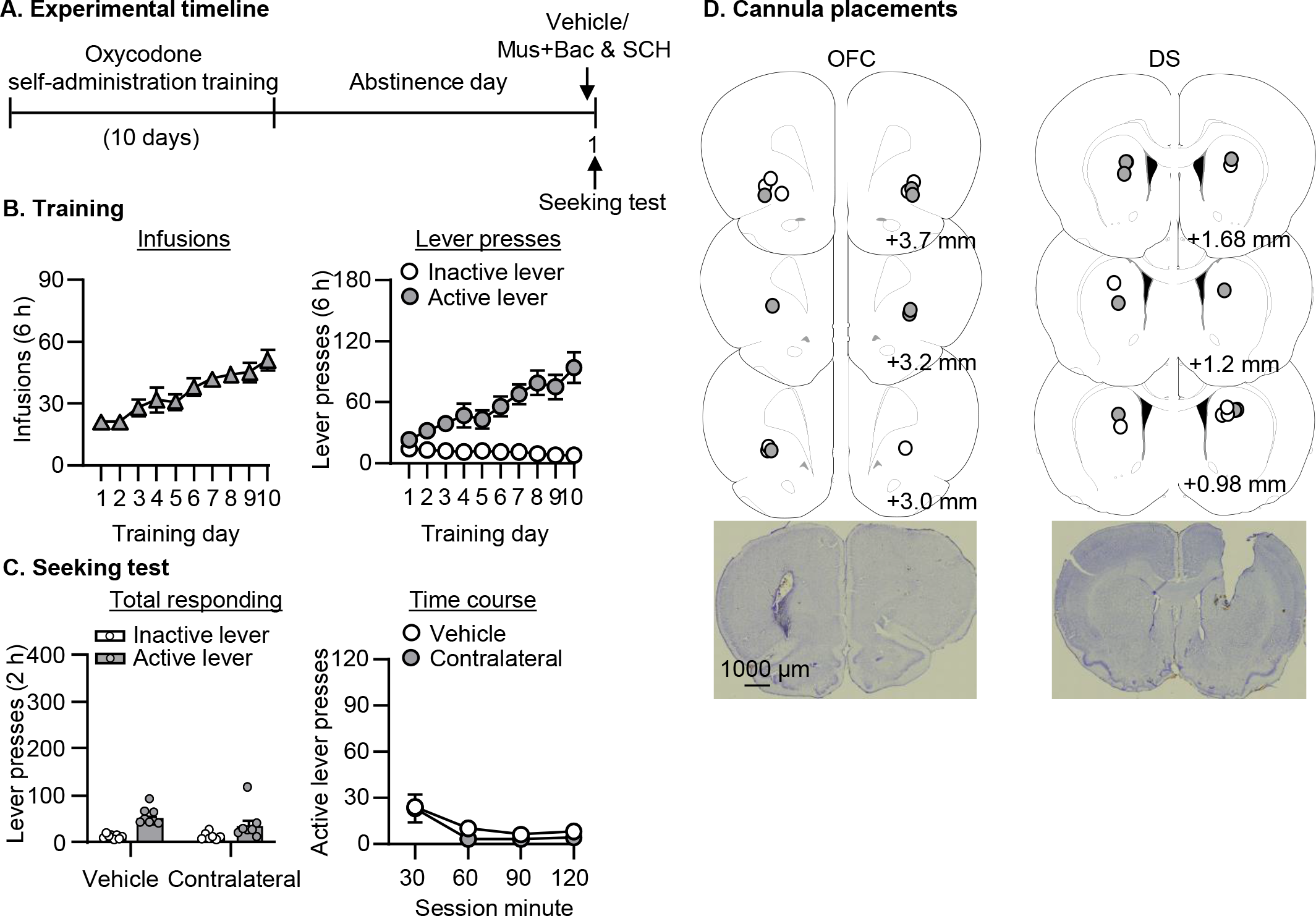
*Contralateral disconnection of OFC or DS had no effect on oxycodone seeking on abstinence day 1 (Experiment 6).* **(A)** Timeline of the experiment. **(B-C)** Oxycodone self-administration. Data are mean ± SEM number of oxycodone (0.1 mg/kg/infusion) infusions or lever presses during the 10 6-h daily self- administration sessions (total n = 15). **(D-E)** Seeking test. Data are mean ± SEM of lever presses on the previously active lever and on the inactive lever during the seeking test sessions. Vehicle, n = 8; Contralateral, n = 7. **(F-G)** Approximate placement [mm from Bregma (Paxinos and Watson, 2005)] of injection tips (vehicle: open circles; Muscimol+baclofen or SCH23390: closed circles), and representative cannula placements.

Finally, to exclude the possibility that the pharmacological disconnection procedures caused motor deficits, we assessed the effect of contralateral disconnection of ipsilateral disconnection on ongoing food self-administration and found neither manipulation impaired ongoing food self-administration. For the contralateral disconnection experiment, the number of food pellets (mean ± SEM) during the 1-h test session was 116±9 after vehicle injections and 110±17 after contralateral disconnection (t(4)=0.408, p=0.70). For the ipsilateral disconnection experiment, the number of food pellets (mean ± SEM) during the 1-h test session was 130±6 after vehicle injections and 123±8 after ipsilateral disconnection (t(4)=0.884, p=0.427).

## Discussion

We investigated the causal role of OFC**→**DS projections in incubation of oxycodone craving. First, we found that DS exhibited time-dependent increases of Fos expression associated with oxycodone seeking after abstinence, and blocking Fos induction by SCH23390, a D1R antagonist, in DS decreased oxycodone seeking on abstinence day 15. Incubated oxycodone seeking was also associated with increased activation of OFC**→**DS projections, indicated by an increased number and percentage of CTb+Fos cells in OFC. Moreover, we found that asymmetrical disconnection of OFC (unilateral injection of muscimol+balclofen into OFC) and DS (unilateral injection of SCH23390 into contralateral or ipsilateral DS), but not unilateral inactivation of OFC or D1R blockade of DS alone, decreased oxycodone seeking on abstinence day 15. Finally, contralateral disconnection of OFC and DS had no effect on oxycodone seeking on abstinence day 1. Together, these data support that the interaction between the activated glutamatergic projections from OFC and D1R signaling in DS is critical for incubation of oxycodone craving.

### Role of dorsal striatum in incubated oxycodone seeking

Our finding that D1R blockade in DS decreases incubated oxycodone seeking extended previous literature on the role of DS in drug seeking across drug classes and animal models of relapse (Fuchs et al., 2005; Vanderschuren et al., 2005; Fuchs et al., 2006; Rogers et al., 2008; Bossert et al., 2009; Jeanblanc et al., 2009; Wang et al., 2010; Pacchioni et al., 2011; Murray et al., 2012; Gao et al., 2013; Rubio et al., 2015; Wang et al., 2015; Li et al., 2015b; Darcq et al., 2016; Caprioli et al., 2017). It is noted that the effect observed here was not due to motor deficits because our previous study found no effect of SCH23390 injections into DS at the same dose on ongoing food self-administration (Li et al., 2015b). One limitation of our current study is that we did not examine the role of D1R signaling in DLS in oxycodone seeking. Several studies have previously documented that DMS and DLS, two DS subregions, play distinct roles in drug-seeking behaviors (Vanderschuren et al., 2005; Bossert et al., 2009; Wang et al., 2010; Corbit et al., 2012; Rubio et al., 2015; Caprioli et al., 2017). In contrast, our previous work found D1R blockade in either DLS or DMS decreased incubated methamphetamine seeking (Li et al., 2015b). Here we observed that oxycodone seeking on either abstinence day 1 or 15 was associated with more Fos expression in DMS than in DLS, suggesting that DMS may play a more prominent role in oxycodone seeking than DLS. Nevertheless, whether D1R signaling in DLS is critical for incubated oxycodone seeking needs to be addressed in future studies.

### Role of OFC**→**DS projections in incubation of oxycodone craving

The main findings in our studies are that asymmetrical disconnections of OFC and DS decreased incubated oxycodone seeking on abstinence day 15. Importantly, the same manipulations had no effect on ongoing food self-administration, which excluded the possibility that the effects observed in the oxycodone studies were due to motor deficits. It is noted that the classical disconnection design used in our study assumes that one brain region transfers information serially to its downstream target in both hemispheres and that learned behaviors can be maintained by one intact hemisphere (Everitt et al., 1991; Gaffan et al., 1993; Floresco et al., 1997; Setlow et al., 2002; Floresco and Ghods-Sharifi, 2007; Bossert et al., 2012). Therefore, the critical role of a pathway in learned behaviors is supported by disrupted behaviors after inactivating the brain region where the pathway originates and the contralateral brain region where the pathway terminates (contralateral disconnection). Furthermore, learned behaviors remain undisturbed after the ipsilateral inactivation of both brain regions in a circuit (ipsilateral disconnection). However, we observed that both contralateral and ipsilateral disconnection of OFC and DS decreased incubated oxycodone seeking. Why did ipsilateral disconnection also inhibit incubated oxycodone seeking?

One possibility is that neural activity in unilateral OFC or D1R signaling in unilateral DS is critical for incubated oxycodone seeking. However, this is unlikely because we found that neither unilateral OFC inactivation nor unilateral dorsal striatal D1R blockade had effects on incubated oxycodone seeking. A second possibility is that incubated oxycodone seeking relies on bilateral OFC**→**DS projections. Supporting this possibility, we found that incubated oxycodone seeking was associated with the comparable activation of both ipsilateral and contralateral OFC**→**DS projections (indicated by a similar number or percentage of CTb+Fos cells in ipsilateral and contralateral OFC). Indeed, previous studies that used this disconnection approach to study corticostriatal circuits with bilateral projections also observed disrupted behaviors after both contralateral and ipsilateral disconnection (Bossert et al., 2012; Jenni et al., 2017; Piantadosi et al., 2020; Jenni et al., 2022). Another interpretation is that activation of ipsilateral OFC**→**DS projections in both hemispheres, instead of one hemisphere, is necessary to maintain incubated oxycodone seeking, which may also responsible for prior studies on the similar inhibitory effect on drug seeking after ipsilateral disconnection (Fuchs et al., 2007; Peters et al., 2008; Lasseter et al., 2011; Bossert et al., 2012; Bossert et al., 2016; Li et al., 2018). To test these possibilities, future studies will use chemogenetic approaches (Roth, 2016; Swanson et al., 2022) to inhibit the OFC**→**DS terminal activities.

In addition, our disconnection data are in line with previous literature implicating OFC**→**DS circuit in compulsive seeking behavior (Ahmari et al., 2013; Burguiere et al., 2013; Gremel et al., 2016; Pascoli et al., 2018). Our findings are also consistent with previous studies demonstrating disrupted incentive motivation to cocaine after contralateral disconnecting OFC and DS projections (Minogianis et al., 2019), increased OFC**→**DS synaptic strengths after the acquisition of cocaine self-administration (Pascoli et al., 2023), and increased OFC-DS circuit strength after methamphetamine self-administration (Hu et al., 2019). However, a rat fMRI study recently reported a negative correlation between OFC-DS functional connectivity and incubation of oxycodone craving after electrical-barrier-imposed voluntary abstinence (Fredriksson et al., 2021). This discrepant observation is most likely due to distinct neural mechanisms underlying forced abstinence (used in the current study) and voluntary abstinence (Venniro et al., 2016; Venniro et al., 2020). Moreover, direct comparison across these two studies is not possible because of the unclear correlation between functional connectivity (assessed by fMRI) and projection activation (assessed by CTb+Fos double labeling), and different time points used in each study (one day after seeking tests for fMRI analysis vs. immediately after seeking tests for CTb+Fos analysis). It is also noted that our findings are conflicted with the role OFC**→**DS in alcohol-related behaviors. For example, withdrawal from chronic intermittent ethanol exposure is associated with decreased excitability of DS-projecting OFC neurons and OFC**→**DS transmissions (Renteria et al., 2018). Consistently, optogenetic induction of long-term potentiation (LTP) of OFC**→**DMS projections decreased alcohol self-administration (Cheng et al., 2021). These findings with alcohol highlight the distinct roles of OFC**→**DS circuits across drug classes.

One limitation here is that we did not examine whether the role of OFC**→**DS in oxycodone seeking relies only on D1R signaling, but not D2R signaling or both, in DS. However, a recent study showed selective potentiation of glutamate transmission from OFC onto D1R- or D2R-expressing medium spiny neurons (MSNs) in DS associated with the acquisition or compulsion of cocaine self-administration (Pascoli et al., 2023). Chronic alcohol exposure also selectively reduces OFC-DS transmission onto D1R-expressing MSNs (Renteria et al., 2018), and the effect of LTP of OFC**→**DMS projections on alcohol self-administration described above depends on D1R, but not D2R signaling in DMS (Cheng et al., 2021). Therefore, we speculate the distinct roles of glutamate transmission from OFC to D1R- versus D2R expressing MSNs in incubation of oxycodone craving, and this question will be examined in future studies.

Finally, it is still unknown how glutamate transmission interacts with postsynaptic D1Rs to mediate incubation of oxycodone craving. One possibility is through striatal cholinergic interneurons (ChIs), which have been major players in mediating striatal dopamine transmission (Cragg, 2006; Cachope and Cheer, 2014; Kosillo et al., 2016; Yorgason et al., 2017). Further support for this hypothesis comes from recent studies indicating that ChIs in striatum contribute to reward seeking(Collins et al., 2019; Cover et al., 2019), cocaine conditioning and extinction (Witten et al., 2010; Lee et al., 2016; Fleming et al., 2022), nicotine seeking(Leyrer-Jackson et al., 2021) and cognitive flexibility after cocaine exposure(Gangal et al., 2023). Another possibility is through retrograde signaling mediated by endocannabinoids. Supporting this hypothesis, previous studies have shown that deletion of cannabinoid type 1 receptor in OFC**→**DS projections disrupts the shift from goal-directed to habitual behaviors in mice (Gremel et al., 2016), and increased retrograde signaling mediated by endocannabinoid from OFC to D1R-expressing MSNs contributes to decreased OFC**→**DS transmission after chronic alcohol exposure (Renteria et al., 2021).

## Concluding remarks

Using neuroanatomical and pharmacological approach, we demonstrated that the interaction between activation of glutamatergic inputs from OFC and D1R signaling in DS plays a critical role in incubation of oxycodone craving after forced abstinence. Our finding extended previous work on the critical role of this circuit in both compulsive-seeking (Ahmari et al., 2013; Burguiere et al., 2013; Gremel et al., 2016; Pascoli et al., 2018) and drug-associated behavior (Renteria et al., 2018; Hu et al., 2019; Minogianis et al., 2019; Wall et al., 2019; Cheng et al., 2021; Fredriksson et al., 2021; Renteria et al., 2021; Pascoli et al., 2023). Interesting questions for future studies include cell-type specificity of dorsal striatal MSNs receiving inputs from OFC and the role of striatal ChIs or endocannabinoid signaling in mediating OFC-DS transmissions. Finally, we used only male rats in the current study, and future studies will examine whether the role of OFC**→**DS in the incubation of oxycodone craving generalizes to female rats, similar to the generalized role of OFC alone in oxycodone relapse across sexes (Altshuler et al., 2021b; Olaniran et al., 2023).

## Acknowledgement

This research was supported by Summer Research Fellowship from University of Maryland College Park (HL), and departmental startup funds (Xuan L). The authors declare that they do not have any conflicts of interest (financial or otherwise) related to the text of the paper. We thank Michael DellaFera for providing the technical support.

## Author contributions

Xuan L conceived the project, provided intellectual inputs, carried out experiments, analyzed the data, and wrote the paper. HL provided intellectual inputs, carried out experiments, analyzed the data, and wrote the paper. AO, Xiang L, JS, MB and CLM carried out experiments.

