## Supplemental Tables 1 and 2 for "Orbitofrontal cortex to dorsal striatum circuit is critical for incubation of oxycodone craving after forced abstinence"

4-9-24

Research article

**after forced abstinence**

Hongyu Lin, Adedayo Olaniran, Xiang Luo, Jessica Strauch, Megan AM Burke, Chloe Matheson and Xuan Li

**Table S1. Statistical analysis**

| Figure number | Test | F value | P-value | Partial ETA^2^ |
| --- | --- | --- | --- | --- |
| Fig. 1A  Exp. 1 DS Fos expression | 3-way ANOVA  Abstinence day * Test condition * Subregion  Subregion: DMS  Abstinence day (between)  Test Condition (between)  Abstinence day * Condition  Subregion : DCS  Abstinence day (between)  Test condition (between)  Abstinence day * Condition  Subregion : DLS  Abstinence day (between)  Test condition (between)  Abstinence day * Condition  Test condition: No test  Abstinence day (between)  Subregion (Within)  Abstinence day * Subregion  Test condition: Seeking test  Abstinence day (between)  Subregion (Within)  Abstinence day * Subregion | F_2,38_ = 5.415  F_1,19_ = 6.377  F_1,19_ = 15.528  F_1,19_ = 9.066  F_1,19_ = 7.724  F_1,19_ = 9.391  F_1,19_ = 8.104  F_1,19_ = 15.109  F_1,19_ = 17.698  F_1,19_ = 18.241  F_1,9_ = 2.782  F_2,18_ = 2.538  F_2,18_ = 2.392  F_1,10_ = 11.202  F_2,20_ = 14.606  F_2,20_ = 4.979 | 0.009*  0.021*  <0.001*  0.007*  0.012*  0.006*  0.01*  <0.001*  <0.001*  <0.001*  0.13  0.107  0.12  0.007*  0.001*  0.018* | 0.222  0.251  0.45  0.323  0.289  0.331  0.299  0.443  0.482  0.49  0.236  0.22  0.21  0.528  0.594  0.332 |
| Fig. 2B  Exp. 2 Bilateral DS inactivation: training | Infusions  Training day (within)  Lever presses  Training day (within)  Lever (within)  Training day * Lever | F_9,81_ = 23.775  F_9,81_ = 8.657  F_1,9_  = 105.776  F_9,81_  = 15.902 | <0.001*  <0.001*  <0.001*  <0.001^*^ | 0.725  0.490  0.922  0.639 |
| Fig. 2C  Exp. 2 Bilateral DS inactivation: seeking test on abstinence day 15 | Total responding (2 h)  Lever (within)  Treatment (between)  Lever * Treatment  Time course  *Active Lever*  Session minute (within)  Treatment (between)  Session minute * Treatment | F_1,8_ = 59.367  F_1,8_ = 8.387  F_1,8_ = 9.528  F_3,24_ = 38.914  F_1,8_ = 9.029  F_3,24_ = 4.477 | <0.001*  0.02*  0.015^*^  <0.001*  0.017^*^  0.012^*^ | 0.881  0.512  0.544  0.829  0.530  0.359 |
| Fig. 3C  Exp. 3 Unilateral CTb injections into DS: training | Infusions  Training day (within)  Lever presses  Training day (within)  Lever (within)  Training day * Lever | F_9,90_ = 7.031  F_9,90_ = 5.888  F_1,10_ = 240.638  F_9,90_  = 5.357 | <0.001*  <0.001*  <0.001*  <0.001* | 0.413  0.371  0.960  0.349 |
| Fig. 3D  Exp. 3 Unilateral CTb injections into DS: seeking test on abstinence day 1 and 15 | Total responding (2 h)  Lever (within)  Abstinence day (between)  Lever * Abstinence day  Time course  *Active Lever*  Session minutes (within)  Abstinence (between)  Session minutes * Abstinence | F_1,9_ = 22.742  F_1,9_ = 15.071  F_1,9_ = 12.128  F_3,27_ = 8.953  F_1,9_ = 14.074  F_3,27_ = 1.732 | 0.001^*^  0.004  0.007^*^  <0.001^*^  0.005^*^  0.184 | 0.716  0.626  0.574  0.499  0.61  0.161 |
| Fig. 3E-G  Exp. 3 Unilateral CTb injections into DS: number of CTb+Fos in OFC on abstinence day 15 | CTb  Hemisphere (Within)  Group (Between)  Hemisphere * Group  Fos+CTb  Hemisphere (Within)  Group (Between)  Hemisphere * Group  Fos+CTb%  Hemisphere (Within)  Group (Between)  Hemisphere * Group | F_1,9_ = 2.639  F_1,9_ = 3.06  F_1,9_ = 1.397  F_1,9_ = 0.973  F_1,9_ = 36.734  F_1,9_ = 3.94  F_1,9_ = 1.964  F_1,9_ = 118.702  F_1,9_ = 0.001 | 0.139  0.114  0.268  0.35  <0.001*  0.078  0.195  <0.001*  0.97 | 0.227  0.254  0.134  0.098  0.803  0.304  0.179  0.93  0 |
| Fig. 4B  Exp. 4 Contralateral disconnection of OFC🡪DS projections: training | Infusions  Training day (within)  Lever presses  Training day (within)  Lever (within)  Training day * Lever | F_9,153_ = 18.221  F_9,153_ = 2.612  F_1,17_ = 181.922  F_9,153_ = 25.855 | <0.001*  <0.001*  <0.001*  <0.001* | 0.517  0.133  0.915  0.603 |
| Fig. 4C  Exp. 4 Ipsilateral disconnection of OFC🡪DS projections: training | Infusions  Training day (within)  Lever presses  Training day (within)  Lever (within)  Training day * Lever | F_9,108_ = 5.861  F_9,108_ = 7.733  F_1,12_ = 89.665  F_9,108_ = 12.167 | <0.001*  <0.001*  <0.001*  <0.001* | 0.328  0.392  0.882  0.503 |
| Fig. 4D  Exp. 4 Contralateral disconnection of OFC🡪DS projections: seeking test on abstinence day 15 | Total responding (2 h)  Lever (within)  Treatment (between)  Lever * Treatment  Time course  *Active Lever*  Session minutes (within)  Treatment (between)  Session minutes * Treatment | F_1,16_ = 99.483  F_1,16_ = 7.126  F_1,16_ = 9.511  F_3,48_ = 53.141  F_1,16_ = 8.362  F_3,48_ = 0.765 | <0.001*  0.017^*^  0.007^*^  <0.001*  0.011^*^  0.52 | 0.861  0.308  0.373  0.769  0.343  0.046 |
| Fig. 4D  Exp. 4 Ipsilateral disconnection of OFC🡪DS projections: seeking test on abstinence day 15 | Total responding (2 h)  Lever (within)  Treatment (between)  Lever * Treatment  Time course  *Active Lever*  Session minutes (within)  Treatment (between)  Session minutes * Treatment | F_1,11_ = 36.638  F_1,10_ = 4.851  F_1,11_ = 6.248  F_3,33_ = 23.064  F_1,11_ = 5.594  F_3,33_ = 3.62 | <0.001*  0.05^*^  0.03*  <0.001*  0.037*  0.023* | 0.769  0.306  0.362  0.677  0.337  0.248 |
| Fig. 5B  Exp. 5  Unilateral OFC inactivation : training | Infusions  Training day (within)  Lever presses  Training day (within)  Lever (within)  Training day * Lever | F_9,126_ = 13.206  F_9,126_ = 4.823  F_1,14_ = 74.357  F_9,126_ = 11.992 | <0.001*  <0.001*  <0.001*  <0.001* | 0.485  0.256  0.842  0.461 |
| Fig. 5C  Exp. 5  Unilateral DS inactivation : training | Infusions  Training day (within)  Lever presses  Training day (within)  Lever (within)  Training day * Lever | F_9,135_ = 29.157  F_9,135_ = 12.447  F_1,15_ = 72.663  F_9,135_ = 17.796 | <0.001*  <0.001*  <0.001*  <0.001* | 0.66  0.454  0.829  0.543 |
| Fig. 5B  Exp. 5  Unilateral OFC inactivation : seeking test on abstinence day 15 | Total responding (2 h)  Lever (within)  Treatment (between)  Lever * Treatment  Time course  *Active Lever*  Session minutes (within)  Treatment (between)  Session minutes * Treatment | F_1,13_ = 70.485  F_1,13_ = 0.408  F_1,13_ = 0  F_3,39_ = 34.485  F_1,13_ = 0.133  F_3,39_ = 0.257 | <0.001*  0.534  0.995  <0.001*  0.721  0.856 | 0.844  0.03  0  0.726  0.01  0.019 |
| Fig. 5B  Exp.5  Unilateral DS inactivation : seeking test on abstinence day 15 | Total responding (2 h)  Lever (within)  Treatment (between)  Lever * Treatment  Time course  *Active Lever*  Session minutes (within)  Treatment (between)  Session minutes * Treatment | F_1,14_ = 27.735  F_1,14_ = 0.117  F_1,14_ = 0.047  F_3,42_ = 32.689  F_1,14_ = 0.079  F_3,42_ = 0.078 | <0.001*  0.737  0.832  <0.001*  0.783  0.971 | 0.608  0.008  0.003  0.7  0.006  0.006 |
| Fig. 6B  Exp. 6 Contralateral disconnection of OFC🡪DS projections: training | Infusions  Training day (within)  Lever presses  Training day (within)  Lever (within)  Training day * Lever | F_9,126_ = 15.961  F_9,126_ = 6.193  F_1,14_ = 60.601  F_9,126_ = 9.194 | <0.001*  <0.001*  <0.001*  <0.001* | 0.533  0.307  0.812  0.396 |
| Fig. 6C  Exp. 6 Contralateral disconnection of OFC🡪DS projections: seeking test on abstinence day 1 | Total responding (2 h)  Lever (within)  Treatment (between)  Lever * Treatment  Time course  *Active Lever*  Session minutes (within)  Treatment (between)  Session minutes * Treatment | F_1,13_ = 27.377  F_1,13_ = 0.974  F_1,13_ = 1.626  F_3,39_ = 15.004  F_1,13_ = 1.302  F_3,39_ = 0.295 | <0.001*  0.342  0.225  <0.001*  0.275  0.829 | 0.678  0.07  0.111  0.536  0.091  0.022 |

**Table S2. Post-hoc analysis**

| Figure Number | Factor Name | Mean Difference | Std. Error | Adjusted p-value | 95% confidence interval  (lower bound to upper bound) |
| --- | --- | --- | --- | --- | --- |
| Fig. 1A  Exp. 1 DS Fos expression | Seeking Test : AD1 vs. AD15  DMS  DCS  DLS  No test : AD1 vs. AD15  DMS  DCS  DLS  AD1 : Seeking test  DMS vs. DCS  DMS vs. DLS  DCS vs. DLS  AD15 : Seeking test  DMS vs. DCS  DMS vs. DLS  DCS vs. DLS | -66.5  -17  -36.167  5.833  0.8  0.433  15.167  16.5  -1.33  45.5  66  20.5 | 22.611  3.952  12.224  3.649  0.338  0.405  5.288  5.926  0.76  12.428  20.801  9.992 | 0.015*  0.002*  0.014*  0.144  0.042*  0.313  0.035*  0.039*  0.14  0.015*  0.025*  0.095 | -116.881 to -16.119  -25.807 to -8.193  -63.404 to -8.93  -2.421 to 14.088  0.036 to 1.564  -0.484 to 1.35  1.574 to 28.759  1.267 to 37.733  -3.287 to 0.621  13.553 to 77.447  12.53 to 119.47  -5.187 to 46.187 |
| Fig.2C  Exp. 2 Bilateral DS inactivation: seeking test on abstinence day 15 | Lever  Active lever  Time course (active lever)  30 minutes  60 minutes  90 minutes  120 minutes | 115.40  48.4  28.4  26.2  12.4 | 38.404  14.858  10.193  11.207  8.118 | 0.017*  0.012*  0.024*  0.048*  0.192 | 26.84 to 203.96  14.137 to 82.663  4.895 to 51.905  0.356 to 52.044  -6.32 to 31.12 |
| Fig. 3D  Exp. 3 Unilateral CTb injections into DS: seeking test on abstinence day 1 and 15 | Lever  Inactive lever  Active lever  Time course (active lever)  30 minutes  60 minutes  90 minutes  120 minutes | -19.964  -133.679  -48.857  -38  -28.964  -17.857 | 6.822  35.633  19.855  9.455  9.311  6.938 | 0.017*  0.005*  0.036*  0.003*  0.013*  0.03* | -35.397 to -4.532  -214.287 to -53.07  -93.771 to -3.943  -59.389 to -16.611  -50.028 to -7.901  -33.553 to -2.162 |
| Fig. 3E-G  Exp. 3 Unilateral CTb injections into DS: number of CTb+Fos in OFC on abstinence day 15 | Fos+CTb  Ipsi : No-test vs Seeking test  Contra : No-test vs Seeking test  Fos+CTb%  Ipsi : No-test vs Seeking test  Contra : No-test vs Seeking test | -127  -98.179  -0.599  -0.604 | 23.731  15.245  0.055  0.102 | <0.001*  <0.001*  <0.001*  <0.001* | -180.684 to -73.361  -132.666 to 63.691  0.723 to -0.475  -0.834 to -0.373 |
| Fig. 4D  Exp. 4 Contralateral disconnection of OFC🡪DS projections: seeking test on abstinence day 15 | Lever  Active lever  Time course (active lever)  30 minutes  60 minutes  90 minutes  120 minutes | 56.625  17.250  15.4  16.625  7.35 | 19.582  9.613  7.67  2.929  4.376 | 0.011*  0.092  0.062  <0.001*  -0.112 | 15.113 to 98.137  -3.13 to 37.63  -0.859 to 31.659  10.417 to 22.833  -1.927 to 16.627 |
| Fig. 4E  Exp. 4 Ipsilateral disconnection of OFC🡪DS projections: seeking test on abstinence day 15 | Lever  Active lever  Time course (active lever)  30 minutes  60 minutes  90 minutes  120 minutes | 96.929  50.167  21.186  17.595  7.381 | 40.983  21.422  11.304  6.264  8.485 | 0.037*  0.039*  0.08  0.017*  0.403 | 6.727 to 187.13  3.017 to 97.316  -3.094 to 46.665  3.808 to 31.383  -11.295 to 26.065 |
